# Morphologically defined substages of tail morphogenesis in *C. elegans* males

**DOI:** 10.1101/2024.01.11.575265

**Authors:** Karin Kiontke, Porfirio Fernandez, Alyssa Woronik, David H. A. Fitch

**Affiliations:** Department of Biology, New York University, 100 Washington Square E., New York, NY 10003; Sacred Heart University, 5151 Park Avenue, Fairfield, CT 06825

**Keywords:** Developmental timing, basement membrane, cell fusion, apical extracellular matrix, nuclear migration

## Abstract

**Background:** Sex-specific morphogenesis occurs in *C. elegans* in the vulva of the hermaphrodite and in the male tail during the last larval stage. Temporal progression of vulva morphogenesis has been described in fine detail. However, a similar precise description of male tail morphogenesis was lacking.

**Results:** We here describe morphogenesis of the male tail at time points matching vulva development with special focus on morphogenesis of the tail tip. Using fluorescent reporters, we follow changes in cell shapes, cell fusions, nuclear migration, modifications in the basement membrane and formation of a new apical extracellular matrix at the end of the tail.

**Conclusion:** Our analysis answers two open questions about tail tip morphogenesis (TTM) by showing that one of the four tail tip cells, hyp11, remains separate while the other cells fuse with each other and with two additional tail cells to form a ventral tail syncytium. This fusion begins early during TTM but is only completed towards the end of the process. This work provides a framework for future investigations of cell-biological factors that drive male tail morphogenesis.

## 1 Introduction

In *C. elegans,* non-gonadal sex differences develop during the last two larval stages before the animals become adults. Development of the vulva in hermaphrodites has been studied as a powerful model for differentiation and morphogenesis for decades (Schindler & Sherwood, 2013). Recently, the progression of vulva morphogenesis during the 4th larval stage was carefully followed and described in 10 morphological substages (Mok et al., 2015). In males the copulatory structures in the tail develop during the 4th larval stage: a pair of spicules, sensilla surrounding the cloaca, and 9 pairs of sensory papillae (rays) that are embedded in a cuticular fan (Emmons, 2005). At the same time, the tail tip changes its shape from long and pointed to short and round in a process called Tail Tip Morphogenesis (TTM). TTM has been studied in some detail at the morphological level (Nguyen et al., 1999; Sulston et al., 1980), and with respect to its regulation (Del Rio-Albrechtsen et al., 2006; Herrera et al., 2016; Kiontke et al., 2019; Mason et al., 2008; Nelson et al., 2011; Sulston et al., 1980; Zhao et al., 2002).

The most detailed morphological descriptions of TTM are found in the papers by Sulston et al. (1980) and Nguyen et al. (1999), which used differential interference contrast microscopy, antibody staining and transmission electron microscopy to observe gross morphological changes, nuclear divisions, changes in nuclear positions and the fate of the adherens junctions (AJs) between tail tip cells during TTM. These studies showed that the tail tip of males and hermaphrodites consists of 4 cells, hyp 8-11, that originate during embryogenesis. In males only, an additional binucleate cell, hyp13, is present ventrally just anterior of the tail tip. In males, at the beginning of the L4 stage, the AJs between hyp8-11 disassemble. Subsequently, the tail tip tissue rounds up and shortens. In the course of this process, the nuclei of the tail tip cells are displaced anteriorly.

Since those early studies, many fluorescent markers for cell biological components have been developed. Some, like those for AJs and non-muscle myosin, were evaluated in the context of TTM (Mason et al., 2008; Nelson et al., 2011) . Observation with fluorescently marked neurons in the tail showed that these also undergo male-specific morphogenesis. In L3 males and hermaphrodites, two PHC neurons send a dendritic process into the tip of the tail (hyp10) and an axon into the preanal ganglion (White et al., 1986). In males only, during L4, the PHC dendritic process shortens during TTM, and the axon grows out anteriorly and differentiates into a male-specific hub neuron that is required for male mating behavior (Serrano-Saiz et al., 2017). A second pair of male specific neurons, PHDL and PHDR, develop de-novo through transdifferentiation and reorganization of the phasmid socket cells PHso1L and PHso1R (Molina-García et al., 2020). However, until recently, no reliable marker for the tail tip cells themselves was known, which would permit precise description of the shape changes of the tail tip cells during TTM and re-evaluate two questions left open by the previous studies of TTM. (1) Do tail tip cells fully fuse and if so, when during TTM? Nguyen et al. (1999) observed with antibody staining and on TEM sections that AJs between the 4 tail tip cells disassemble early during TTM, and apical plasma membranes of adjacent cells became contiguous. Cytoplasm of adjacent cells was also contiguous just under the former AJ sites. However, cell cytoplasms were still mostly separated by membranes. Sulston et al. (1980) observed a ventral hypodermal syncytium with initially 5, then 7 nuclei, indicating that the tail tip cells fuse with the binucleate hyp13 at one point during TTM, but it was unclear when this fusion occurs. (2) What is the fate of hyp11? Fusion of all tail tip cells hyp8-11, as it is e.g. proposed in WormAtlas (Lints & Hall, 2005), poses a topological problem for tail tip cell retraction: Different from all other tail tip cells, hyp11 is positioned dorsolateral of the basement membrane (BM); hyp8-10 and hyp13 are positioned ventrally (Fig. 2). If hyp11 were to fuse with hyp8-10, it would have to breach the BM or somehow move around the BM to the ventral side. If such a breach actually takes place and when during TTM, was unknown.

RNA-seq of tail tip-specific transcriptomes (Kiontke et al., 2024) identified several genes that are specifically expressed in the tail tip cells. We made translational reporters for two of these genes, *grl-26* and *bli-1* to follow the fate of the tail tip cells throughout TTM in the context of reporters for other cellular components like AJs and the BM, and redescribe the process of TTM. We found that tail tip cells fully fuse only late during TTM and that hyp11 is not part of this syncytium but remains a separate cell dorsal of the BM. We find that the zona pellucida proteins NOAH-1 and NOAH-2 are secreted into a new apical ECM at the end of the tail. We provide a description of TTM at a fine temporal resolution matching that for the vulva published by Mok et al. (2015), which enables evaluating the developmental age of individual animals without following their development.

## 2 Results

To match TTM developmental substages to those determined for vulva development, we synchronized *C. elegans* by the hatchoff method, which collects larvae that hatched within one hour. These L1s were grown at 20 or 25 °C, and hermaphrodites and males observed every hour beginning with 32 or 22 h, respectively, after hatch. Vulva substages were identified after Mok et al. (2015). As with vulva development, more than one substage of TTM was observed simultaneously at every collection time point, indicating that there is individual variation in developmental rate. Thus, the times after hatch for the different stages given in Fig. 1 are approximate.

**Figure 1.**
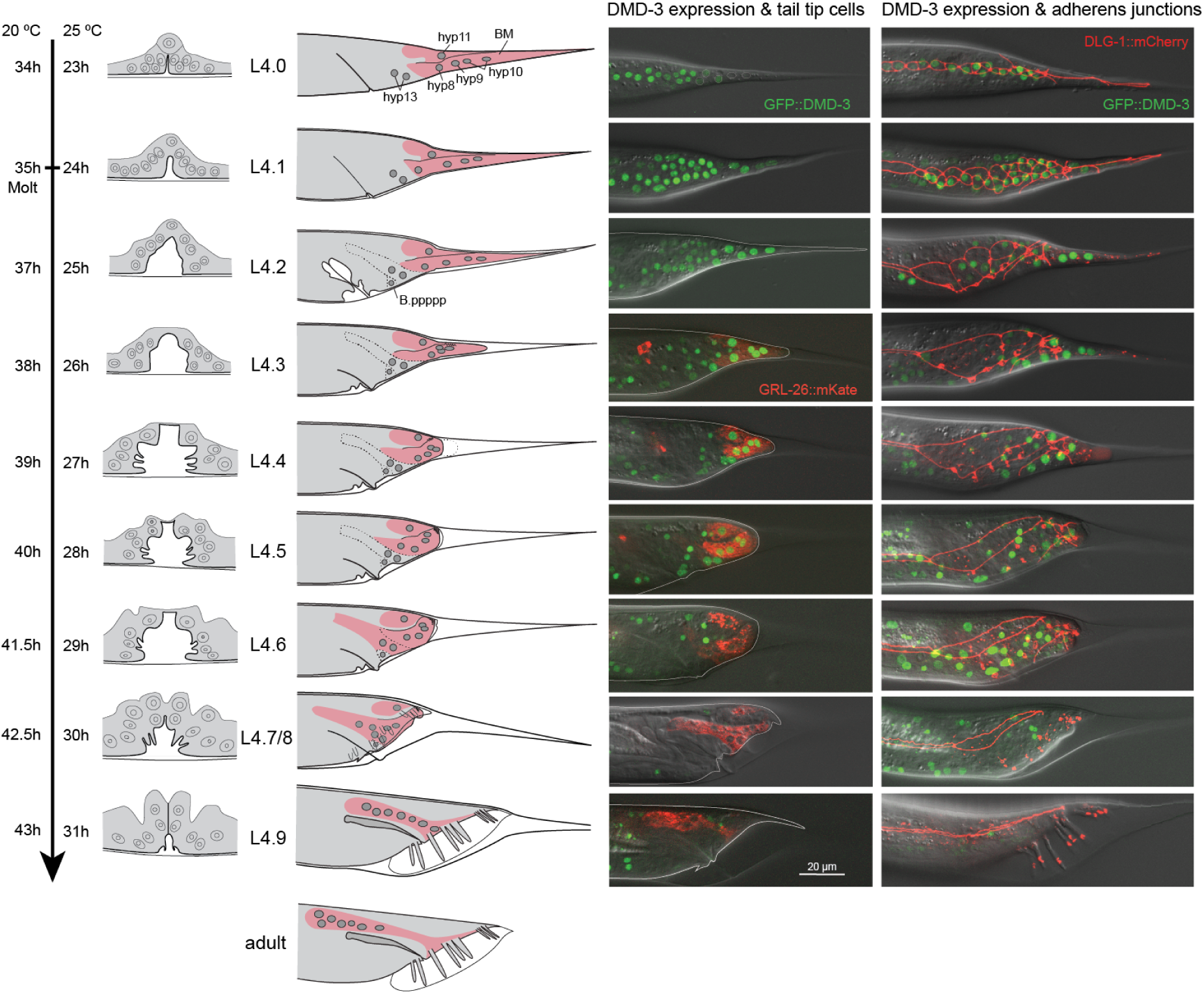
Overview of TTM in male *C. elegans* in comparison to vulva morphogenesis at equivalent time points. First column: schematic reconstruction of the hermaphrodite vulva from L3 to late L4 labeled as in Mok et al. (2015). To the left, the developmental time post-hatch is given for worms grown at 20 °C and 25 °C. Note that there is a considerable spread of stages at each time point; only the most typical morphology for each timepoint is shown. Second column: Schematic reconstruction of the tail tip of males from late L3 to late L4. The nuclei of the tail tip cells are shown as gray circles. The tail tip cells are visualized in pink, dashed lines indicate the cell boundaries of hyp13. Third column: Tails of males expressing endogenous GFP::DMD-3 in nuclei and endogenous GRL-26::mKate in the cytoplasm. Fourth column: Tails of males expressing GFP::DMD-3 and DLG-1::mCherry, which marks adherens junctions.

To study the process of TTM, we observed strains expressing fluorescently marked proteins. We used DLG-1::mCherry to mark AJs (Fig. 1), LAM-1::GFP to mark the BM (Fig. 2), and a construct expressing mCherry fused to the human pleckstrin homology (PH) domain under the *lin-44* promoter to mark the tail tip membranes (Figure 3). *lin-44* is only expressed in tail tip cells from embryogenesis through adulthood [Herman & Horvitz, 1995]. GRL-26::mKate or BLI-1::GFP are expressed in the male tail tip cytoplasm (Fig. 1 and 4). A GFP reporter for the transcription factor DMD-3 is expressed in the tail tip nuclei. We also found that two components of the apical extracellular matrix, the zona pellucida proteins NOAH-1 and NOAH-2, are expressed during TTM. Using these tools and previous knowledge (Nguyen et al., 1999; Sulston et al., 1980), we provide a detailed description of the process of TTM and break it up into 10 stages that match those defined for vulva development by Mok et al. (2015) (Fig. 1).

**Figure 2.**
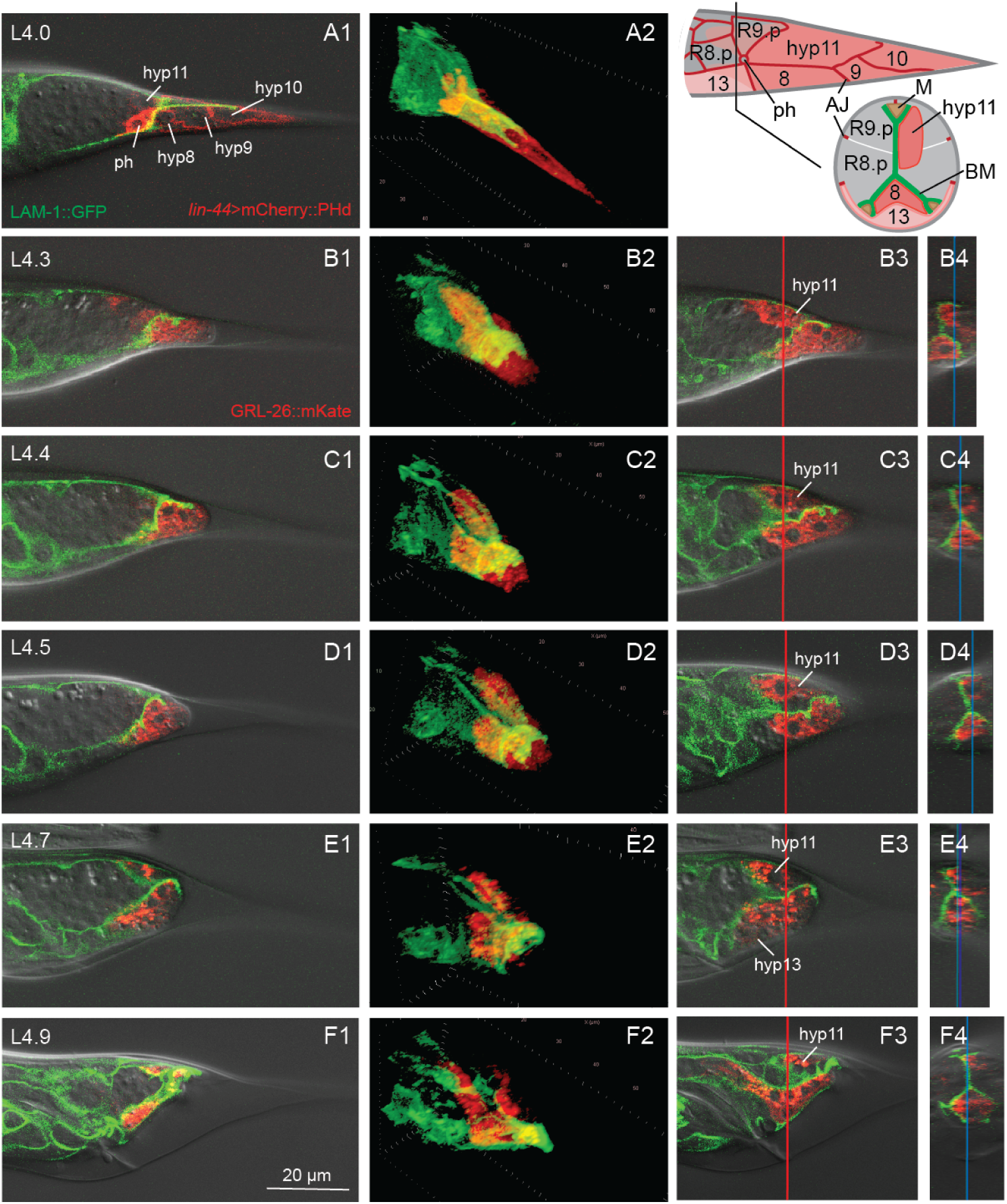
Remodeling of the basement membrane during TTM. Male tails over the course of TTM (A-F) with the BM marked with LAM-1::GFP and the tail tip cells marked with GRL-26::mKate, except A, where the tail tip cell membrane is labeled with *lin-44*>mCherry::PHdomain::*lin-44*-3’UTR. Top right: Schematic of the tail tip surface of a male before the onset of TTM and schematic cross section at a level slightly anterior of the phasmid opening (ph). AJs in red, tail tip cells hyp8-11 in pink, hyp13 in light pink, other epidermal cells in gray, BM in green, muscle cells (M) in brown. The tail tip cells abut R9.p, a cell derived from the ray 9 lineage, and hyp13. hyp8 and hyp11 send anteriorly directed processes into the tail. The process of hyp11 is asymmetrically located to one side of the BM. The BM forms a sheath over hyp8-10 and covers each of two ventral and one dorsal muscle cells (modified after Nguyen et al., 1999).

**Figure 3.**
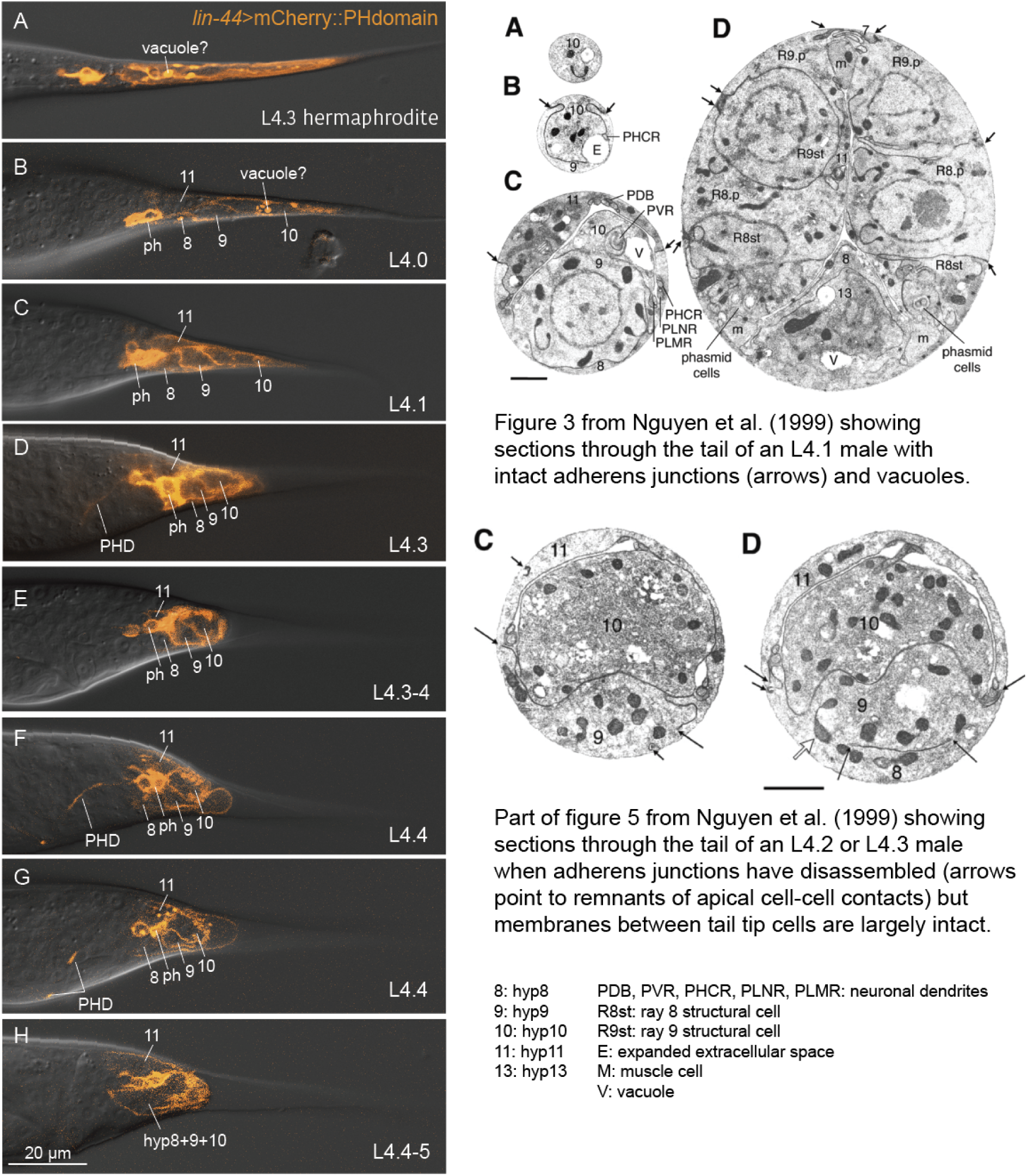
Membranes of tail tip cells disassemble late during TTM. Left panel: A construct expressing mCherry fused to the human PH domain under the promoter of *lin-44* marks the membrane of tail tip cells hyp8-11 and of the phasmid socket cells. In L4.0 and L4.1 males, bright vacuoles are visible in the tail tip cells (A). The membranes between the tail tip cells are visible (C-F) until about stage L4.5 when the reporter becomes diffuse, indicating that the membranes are disassembling (G). Transdifferentiation of the phasmid socket cells PHso1 begins after stage L4.1. The new PHD neuron continues to express the membrane reporter. By stage L4.3-4, the axonal processes of the neurons have reached a position near the anal opening (E and F). Right panel: Sample thin sections published in Nguyen et al. (1999) for comparison. Top: Figure 3A–D posterior to anterior sections from a stage L4.1 male. Short arrows indicate sites of AJs that are still intact at this stage. There is evidence of vacuole production (V) and expansion of extracellular space (E). Bottom: Figure 5 C and D showing sections through a later stage male. Scale bars 1µm.

**Figure 4.**
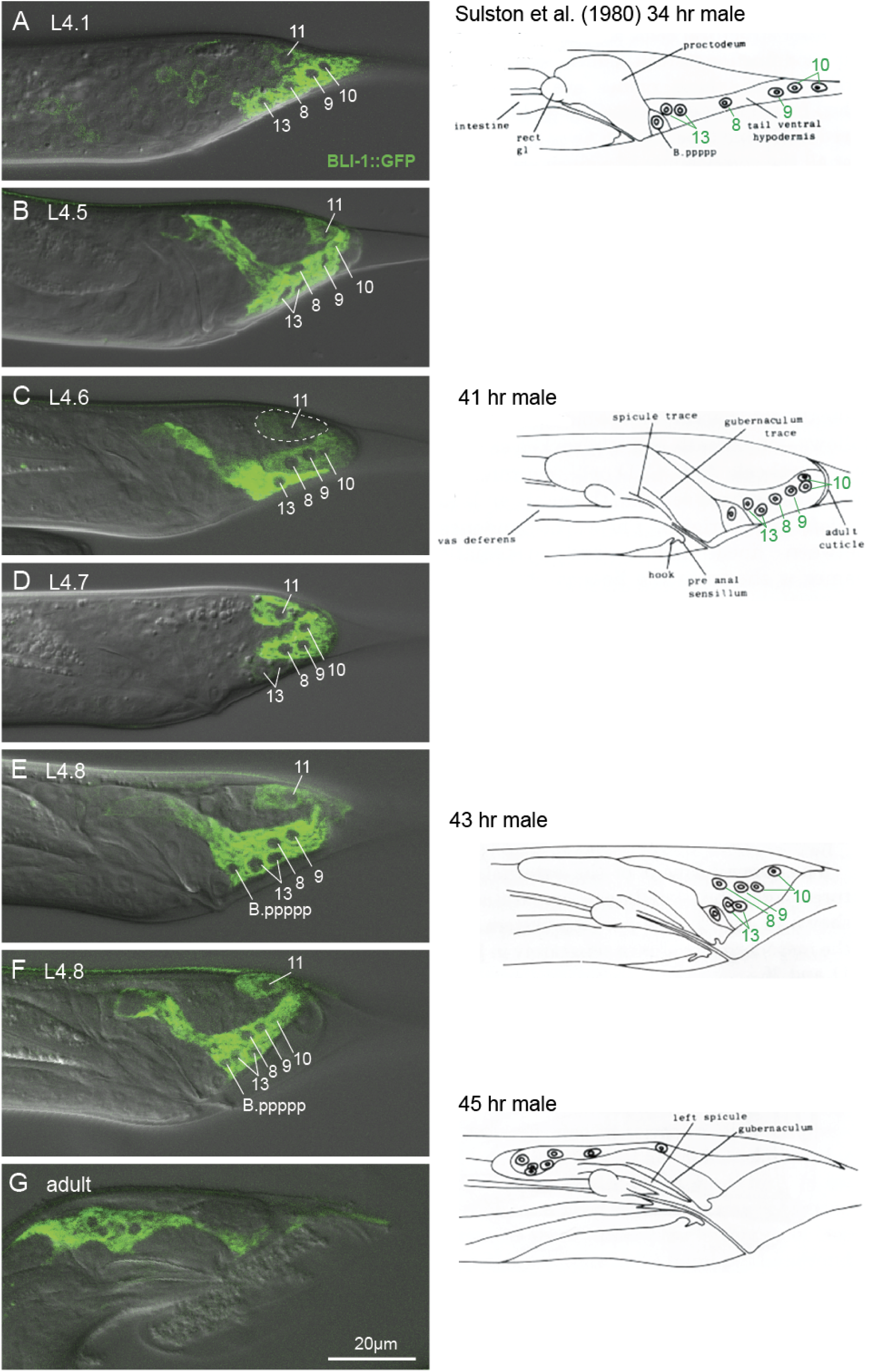
Expression of the multi-copy (extracellular transgene) translational reporter BLI-1::GFP in L4 male tails (left) in comparison with drawings by Sulston et al. (1980). BLI-1::GFP is seen in the cytoplasm of the tail tip cells and hyp13, but not in the nuclei of these cells (labeled). The position of hyp11 is unchanged throughout TTM. All tail tip cells and their fate were identified and interpreted by Sulston et al. (1980). (A) L4.1 male, only one nucleus of hyp10 is in focus, hyp13 is a mononuclear variant. P.ppppp does not express BLI-1. (B) L4.5 male focused on the center of the tail, the anterodorsal process of hyp13 is clearly visible. (C) L4.6 male, hyp8-10 hyp11 and hyp13 are distinguishable by different degrees of reporter expression; the area of hyp11 is outlined for easier identification. (D) In this L4.7 male, hyp13 only weakly expresses BLI-1::GFP, indicating that hyp13 and the tail tip cells are not fully fused. The developmental stage of these animals corresponds to depiction of a male 41 h after hatch by Sulston et al. (1980). The authors did not see a separation of the tail cells and hyp13. (E, F) L4.8 male tails. The reporter is now also expressed in B.ppppp. (G) young adult male, the ventral hyp syncytium and its 7 nuclei have migrated far anteriad. A narrow region of this syncytium remains at the ventral surface of the tail.

Column 1: Z plane focused at the center of the tail tip; Column 2: 3D reconstruction of the same image stack; Column 3: Z plane focused on hyp11; Column 4: Z-Y plane reconstruction at the position indicated by the red line in column 3 images, the blue line indicates the Z plane seen in column 3.

The tail tip cell nuclei were visible either by expression of the transcription factor DMD-3 marked with GFP, or because they exclude the cytoplasmically localized proteins GRL-26::mKate and BLI-1::GFP (Fig. 1 and 4). We quantified the migration of the tail tip nuclei throughout TTM by measuring their distance from phasmid opening as reference point (Fig. 6), as was done previously by Nguyen et al (1999) on ultrastructural data. Anterior migration of the tail tip nuclei begins early during TTM before the tail tip changes shape (between stage L4.1 and L4.2). By mid-L4, all nuclei have reached a position anterior of the phasmid, at which point their migration slows down. Migration accelerates again once the tail tip syncytium has fused with hyp13.

### Stages of male tail development L4.0

Before the L3/L4 molt, the DMD-3 protein is seen in ray-precursor cells and weakly in hyp8, 9 and 11 but not in hyp10. The tail tip cell membranes are clearly visible in strains expressing the lin-44>mCherry::PH reporter. This reporter also marks the phasmid socket cells. The DM-domain transcription factor DMD-3 is expressed in many tail cells during L3, but initially not in the tail tip. At this time, weak expression begins in hyp 8, 9 and 11. DMD-3 is the master-regulator for TTM, and, together with its paralog MAB-3 required for morphogenesis of all male tail structures (Mason et al., 2008). Reporters for GRL-26 and BLI-1, which are under the control of DMD-3 (Kiontke et al., 2024), are not expressed yet. The BM in the tail tip forms a “roof” over hyp8, hyp9 and the anterior part of hyp10. Further anterior, the BM forms a spine-like dorsal process between the Rn.p cells of both sides. hyp11 is located on one side (mostly left) of this spine and dorsal of the roof.

### L4.1 (immediately after the L4/L3 molt)

This stage is equivalent to stage 0 in Nguyen et al. (1999) and to the 34 h male shown by Sulston et al. (1980) (also Fig. S1). The male has completed ecdysis but TTM has not yet begun. DMD-3 expression begins in the hyp10 nuclei. The AJs between the tail tip cells are intact. The membrane marker shows large bright spots inside of the tail tip cells, which we interpret as the vacuoles described by Nguyen et al. (1999) for early L4 males (Fig. 3). These structures are also visible in tail tips of L3 and L4 hermaphrodites (not shown).

#### L4.2

This stage is equivalent to stage 2 in Nguyen et al. (1999). The tail tip cells have dissociated from the L4 cuticle. GFP::DMD-3 is brightly expressed in the nuclei of all tail tip cells and hyp13. The AJs begin disassembly starting in the anterior of the tail tip. Cell membranes between the tail tip cells are present, as well as vacuoles. Reporters for BLI-1 begin to be expressed in the tail tip cells and in hyp13. hyp13, located just posterior of the proctodeum, possesses a central antero-dorsal process abutting the proctodeum. hyp8 abuts hyp13 posteriorly. PHso1 cells are moving away from the phasmid socket and begin the process of transdifferentiation into PHD neurons (Fig. 3, Molina-García et al., 2020).

#### L4.3

The terminus of the tail tip rounds up and begins to shorten. The cytoplasm at the posterior end of the tail is clear. The nuclei of hyp10 migrate anteriad. The anterior hyp10 nucleus can move to a position anterior of the hyp9 nucleus during this stage. All nuclei express GFP::DMD-3. The AJs between the tail tip cells have disassembled with remnants seen in the region of hyp10. There are still a few vacuoles visible inside the tail tip cells.

GRL-26::mKate begins to be expressed in the tail tip cells but not in hyp13. The posterior of the BM begins to shorten and appears scrunched up in accordion fashion. NOAH-2::GFP is seen at the dorsal surface of the tail tip (Fig. 5). The axon of the PHD neuron is forming.

**Figure 5.**
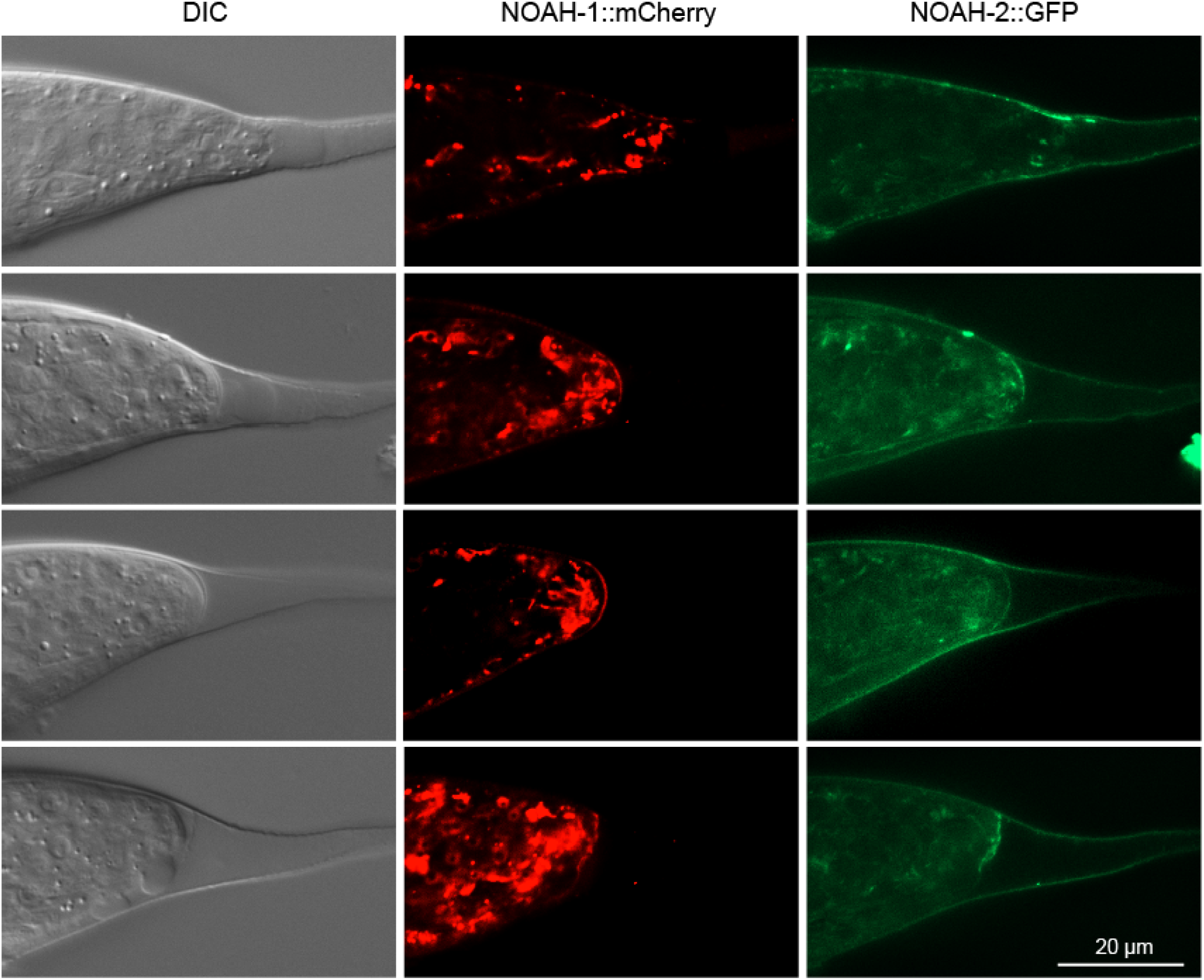
The zona pellucida proteins NOAH-1 and NOAH-2 are deposited in the extracellular matrix that forms a cap at the end of the tail after the tip has rounded up. Images show three separate channels of one Z slice taken of animals that express both reporters.

#### L4.4

The tail tip has rounded up and continues to shorten. At this stage, a membranous “bubble” is often seen in males mounted on a microscope slide. Occasionally, we observed how the bubble forms. We interpret this as an artifact caused by the pressure of the coverslip. At the posterior of the tail, a cap of extracellular matrix is being laid down. It contains NOAH-1 and NOAH-2 (Fig. 5). The bubble appears in places where the cap is either incomplete or breached. All tail tip nuclei and those of hyp13 express DMD-3::GFP brightly. hyp8 sends a short process along the dorsal edge of hyp13. At the beginning of this stage, the membranes between tail tip cells are still visible, but vacuoles are no longer observed (Fig. S4). The BM is further scrunched and shortened. Early during this stage, the axons of the PHD neurons grow out ventrally and anteriorly to a position near the anus. They then grow dorsally and anteriorly.

#### L4.5

The cap of extracellular matrix at the end of the tail is clearly visible with DIC and in animals expressing NOAH-1::mCherry and NOAH-2::GFP. The tail tip cell tissue continues to retract away from this cap. Stage L4.5 corresponds to the 41 h male depicted by Sulston et al. (1980) and possibly stage 3 in Nguyen et al. (1999). The tail tip nuclei are now more difficult to distinguish from other cells in the tail. Nevertheless, Sulston et al. (1980) described the position of hyp8-10 and hyp13 accurately. At the beginning of this stage, the membranes between the tail tip cells disappear and the membrane marker appears diffuse in the cytoplasm (Fig. 3). Further anterior, R1.p-R5.p have fully fused to form the tail seam, which narrows. Ventrally, the tail tissue dissociates from the cuticle in the region posterior to the anus where the cell B.ppppp is located, but remains attached near the now retracted tail tip. Expression of DMD-3 reporters is diminishing in the nuclei of hyp8-11. Animals that express the BLI-1::GFP reporter more or less intensely in hyp13 than in the tail tip cells, as well as the GRL-26 reporter that is only expressed in hyp8-11 show that hyp13 is not fused with the tail tip cells at this stage. hyp11 does not change position. The BM bends ventrally and forms a cap at the end of the tail tip.

#### L4.6

The ventral tail tissue has fully retracted from the L4 cuticle but touches it in the posterior. The end of the tail is squaring off. The tips of the future rays are visible as small buttons. DMD-3 expression is turning off. We assume that hyp8-10 begins fusion with hyp13 and then with B.ppppp to form the ventral tail hypodermis (Fig. 6 L4.6 male middle panel shows faint GRL-26 expression in hyp13 but not in B.ppppp, suggesting that the GRL-26 protein diffuses into hyp13 and that B.ppppp is still separate from this syncytium). Expression of the membrane marker becomes very faint except in the phasmid socket cells and the PHD neuron.

**Figure 6.**
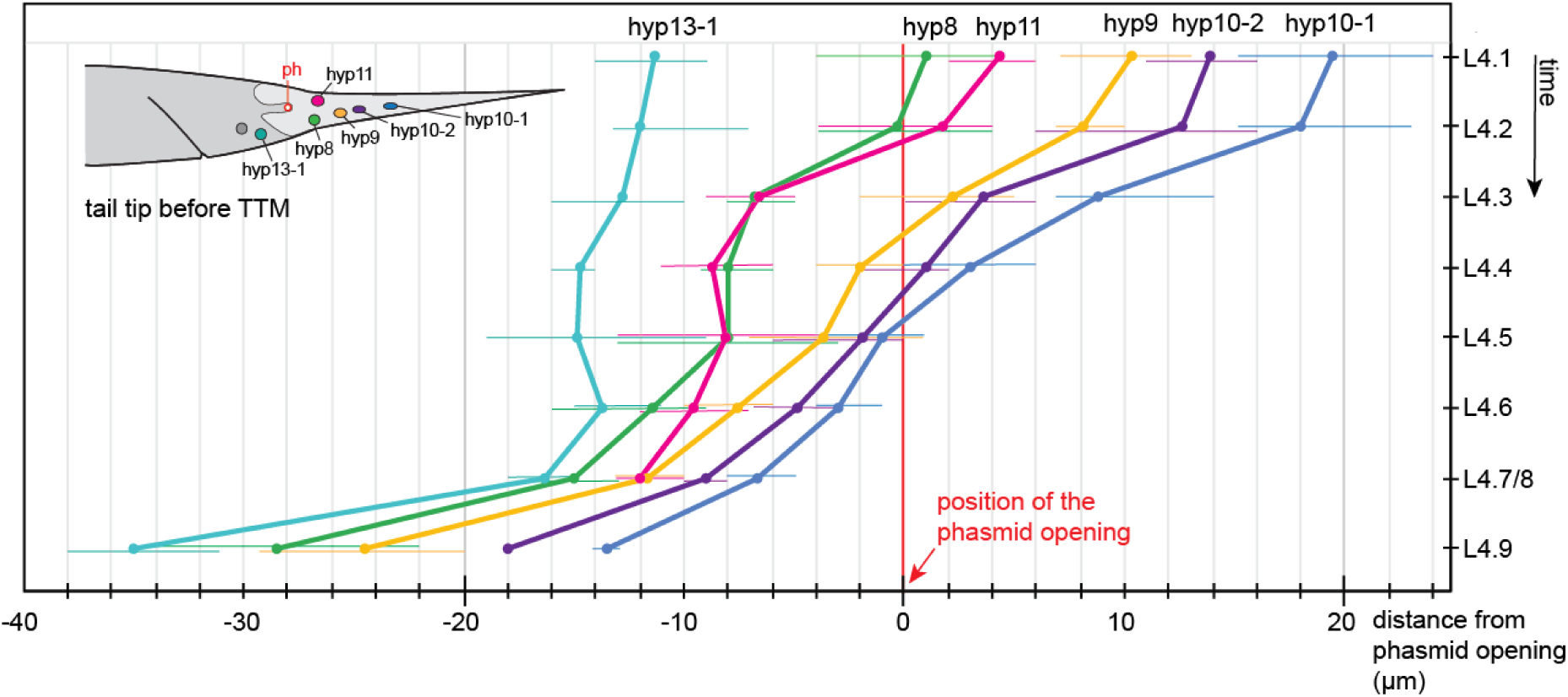
Quantification of the migration of the tail tip nuclei during TTM. Shown is the distance of each nucleus from the location of the phasmid opening as average and measured range (narrow lines). L4.1: n = 9, L4.2: n = 8, L4.3: n = 5, L4.4: n = 7, L4.5: n = 9, L4.6: n = 7, L4.7/8 n = 3, L4.9 n = 2.

**Figure 7.**
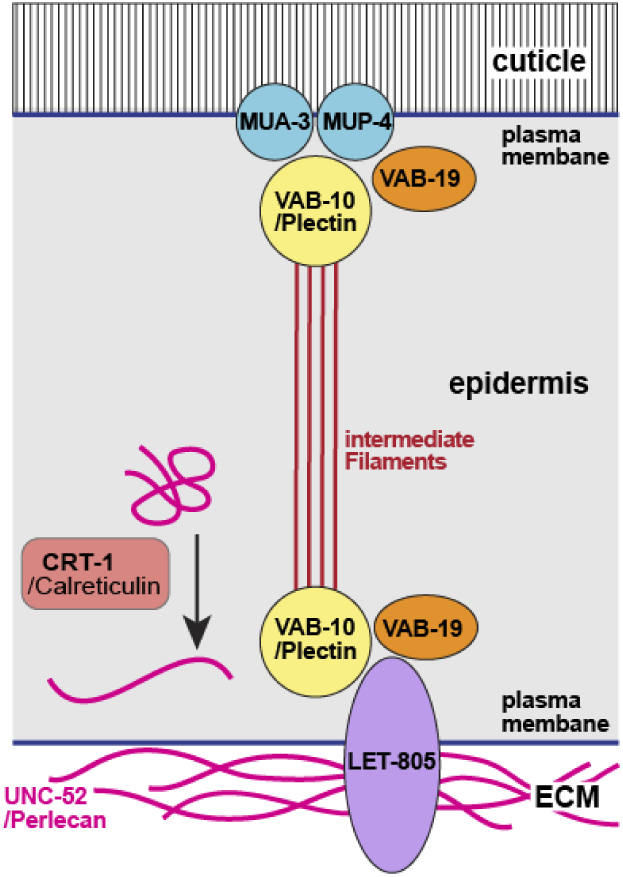
Structure of the *C. elegans* fibrous organelle after Zahreddine et al. (2010)

#### L4.7/8

The ventral tail tissue further retreats antero-dorsally and the rays and fan are being drawn out. During this period, membranous bubbles are often observed at the ventral surface of the tail. The ventral tail hypodermis is fully fused and begins to extend anteriorly. Nuclei of hyp8-10 migrate near the hyp13 nuclei.

#### L4.9 and later

The rays and fan are forming. Anterior migration of the ventral tail hyp continues until after the L4/adult molt and the nuclei of all cells are found in the anterior portion of the syncytium. Expression of GRL-26::mKate and of the BLI-1 reporters in hyp11 turn off before the L4/adult molt. Thus, the location and shape of hyp11 in adults could not be determined.

From our observations we conclude: (1) Even though the AJs between hyp11 and the other tail tip cells disassemble early, hyp11 never fuses with the ventral tail tip cells. During the initial retraction of the tail tip, hyp11 is completely or mostly internalized in a position dorsal to and on one side of the basement membrane. It remains in this position throughout morphogenesis and never breaches the basement membrane or migrates laterally or terminally to the ventral side. Whether the apical connection with the other tail tip cells that is established early during TTM is maintained or severed could not be determined here. It is worth pointing out that hyp11 is derived from a different cell lineage (C lineage) than the other tail tip cells (AB lineage). It thus has a different developmental history and could have a different fate from the other tail tip cells. (2) The membranes between hyp8-10 fully disappear only at the beginning of stage L4.5, at the time the aECM cap is visible. Subsequently, a syncytium is formed of hyp8-10, hyp13 and B.ppppp, where B.ppppp fuses last. Sulston et al. (1980) depicted this syncytium in their 43 h and 45 h male. Consistent with their observation, we find 7 nuclei in this ventral tail hyp: 4 nuclei of hyp8-10, 2 nuclei of hyp13 and one of B.ppppp. (3) Morphogenesis of the anterior part of the tail (dorsoventral flattening and formation of the fan) happens after stage L4.5 and coincides with the formation of the ventral hyp and its anterior migration. (4) The BM in the tail tip is folded accordion-like while the tail tip is shortening. This indicates that the shortening of the BM is passive, i.e. it is pulled by forces in the tail tip cells that are attached to it. The intensity of the LAM-1::GFP in the signal appears to increase in the course of TTM, indicating that the BM is not disassembled but is compacted instead.

## 3 Discussion

The study of fluorescent reporters for various proteins in the tail tip permitted us to redescribe the process of male TTM in further detail than was previously possible. We matched the development of the male tail to that of the vulva. With this guide, the approximate developmental age of an animal at the L4 stage can be estimated without tight synchronization, and dynamic gene expression can be described with precision.

We could answer two open questions about TTM: We found that the tail tip cells hyp8-10 fully fuse with one another only in mid-L4 (stage L4.5) when the tail tip has fully retracted, and subsequently fuse with hyp13 and B.ppppp. The dorsally located hyp11 does not fuse with this ventral syncytium. While the analysis of new reporter genes contributed to our understanding of the process of TTM, many questions about the process are still open including the following:

1. What ensures the integrity of the tail tip cells after they dissociate from the cuticle? Recently, it was shown that a transient apical extracellular matrix (aECM) is required for the integrity of the excretory duct during molts (Gill et al., 2016), and is put down prior to secretion of the body cuticle during each molt, where it helps to shape cuticular structures like alae (Katz et al., 2022). It was speculated that this transient provisional aECM helps maintain the integrity of the body shape of nematodes during the molt (Katz et al., 2022). An aECM at the tip of male tails is clearly discernible after TTM stage L4.4, and at least two proteins of the provisional aECM, NOAH-1 and NOAH-2 are expressed in the tail tip. RNAi against the gene for NOAH-2 causes Lep tails in adult males (Nelson et al., 2011). However the tail tip tissue dissociates from the cuticle much earlier, and the subsequent rapid shortening of the tail tip makes it unlikely that a relatively rigid aECM is stabilizing its shape. Instead, we speculate that a meshwork of intermediate filaments (IF) could support the retracting tail tip cells similar to that in the terminal web under the luminal membrane of the excretory canal (Buechner et al., 2020) until the aECM is laid down. Consistent with this idea, the genes for the IF IFC*-2*, and the IF regulators BBLN-1 (Remmelzwaal et al., 2021) and SMA-5 (MAPKinase) (Geisler et al., 2023) are differentially expressed in tail tips undergoing TTM (Kiontke et al., 2024). Visualization of IFC-2 in the tail tip during TTM can be used to test this hypothesis.
2. How is the cuticle attached to the tail tip cells prior to TTM? Along the nematode body, the cuticle is attached to the epidermis through the so-called fibrous organelles, which consist of apical and basal hemidesmosome-like structures that are connected via IFs (Lažetić & Fay, 2017). The apical hemidesmosome-like structure connects to the cuticle with the EGF repeat proteins MUA-3 and MUP-4. The basal parts of the fibrous organelles connect via LET-805/myotactin to UNC-52/perlecan in the basement membrane and to the underlying muscle cells. The ankyrin-repeat protein VAB-19 is associated with both sides of the fibrous organelle. Abundance of UNC-52 is controlled by CRT-1/Calreticulin (Zahreddine et al., 2010). The tail tip-specific RNA-seq analysis (Kiontke et al., 2024) showed that *mup-4* and *vab-19* are repressed and *crt-1* is activated in tails undergoing TTM. This suggests that fibrous organelles play a role in attaching the cuticle to the tail tip epidermis even in the absence of muscle cells in this part of the body. It is conceivable that the apical hemidesmosomes are released when the tail tip tissue dissociates from the cuticle at the beginning of TTM (hence the downregulation of *mup-4* and *vab-19*). However, the basal hemidesmosomes continue to firmly connect the tail tip cells to the BM. This connection must be very strong, since the BM is scrunched up when the tail tip cells shorten. Upregulation of *crt-1* could play a role in strengthening the BM connection. We thus expect UNC-52 and LET-805 to be expressed at the BM in the tail tip, and mutations in fibrous organelle genes to cause TTM defects.
3. How do the tail tip nuclei move? We and Nguyen et al. (1999) observed the migration of the tail tip nuclei during TTM from a position near the end of the tail in early larvae to one anterior of the cloaca in adults (Fig. 6). How is this migration controlled? It is known that nuclear migration as well as the stability of nuclear positions in syncytia is mediated by the LINC complex, a protein complex that connects the inner and outer nuclear membrane to one another and to microtubules for migration or to the actin cytoskeleton for positional stability (Starr, 2019). We expect that tail tip-specific inactivation of genes in the LINC complex would cause migration defects of tail tip nuclei.

These and other questions about the mechanisms during male tail morphogenesis and its regulation can be addressed by combining fluorescent reporter expression with targeted gene-knockdown either globally by RNAi or by tail-tip specific auxin mediated protein degradation that has recently been optimized for tissue-specific applications in *C. elegans* (Xiao et al., 2023).

## 4 Experimental Procedures

### 4.1 Worm husbandry and synchronization

*C. elegans* strains were grown on a lawn of *E. coli* (OP50-1) according to standard methods (Stiernagle, 2006). To obtain synchronized populations of larvae, we used the hatchoff method, described in Woronik et al. (2022). Briefly, gravid hermaphrodites were allowed to lay eggs overnight. Then, hermaphrodites and all hatched larvae were removed by repeated washes with M9 buffer. The remaining embryos were incubated at 25 °C. After one hour, all L1 that had hatched during this time were collected in M9, placed on food and allowed to develop at 20 °C or 25 °C until the late L3 stage. After this time, several males and hermaphrodites were observed hourly for vulva and male tail development.

### 4.2 Imaging

For microscopy, worms were paralyzed in 20 mM sodium azide and mounted on a pad of 4 % Noble Agar in 50 % M9 buffer supplemented with 20 mM sodium azide. They were examined with a Zeiss AxioImager equipped with Colibri LED illumination and an Apotome. Generally, image stacks were recorded with 0.5 µm distance between slices. If the fluorescent signal was too dim for Apotome imaging, the deconvolution algorithm implemented in the Zeiss ZenBlue software (v. 2.6) was used (“good/fast iterative” setting) to improve image quality. In some cases, several images in a stack were combined in a Z-projection using Zen Blue. 3D reconstructions were also made with Zen Blue.

### 4.3 Measurements

To follow the migration of tail tip nuclei, we measured the distance between the center of each nucleus from the location of the phasmid on images of animals that expressed DLG-1::mCherry to label the adherens junctions at the phasmid opening. The nuclei were either marked with GFP::DMD-3, or were clearly visible in animals expressing cytoplasmic BLI-1::GFP or GRL-26::mKate.

### 4.4 Translational reporter generation

We inserted mKate at the endogenous *grl-26* locus 3’ of the coding region using CRISPR Cas9 genome editing. The algorithms implemented in Benchling (https://www.benchling.com) identified a PAM site 9 nt 3’ of the stop codon. Oligonucleotides with 120 nt long homology arms, a linker and an overhang in the coding sequence for mKate in plasmid pCFJ350 (Addgene) (grl-26+mKate_F ATGGAAATATATAATCGAAATTTTTCCTTTTTTGTAGAATCCCTTCCACTATTCCATCAAACA TGATTCAGCATACTGCGGAGCACGGAATGGTTCTCATTATTGTCAGGCATTTGCGATA*GG AGCATCGGGAGCCTCAGGAGCATCG*atgtccgagctcatcaaggagaacatg; grl-26+mKate_R CAATTGGCTTAAATTCTTAGTCGATTTATTACCGTTTTGCGATGTTAAAGAGCTTGTTATTTATTTCTATCATGAAAATTTATTGGATAATTAAAAACAAAATAGCgTTTCGTAAAGTTAacggtgtccgagcttggatgg; homology arm capital letters, linker italics) were used to PCR amplify (Takara PrimeStar Max) the homology repair template (HRT). After purification (Promega Wizard SV Gel and PCR clean-up system), the concentration of the HRT was adjusted to 100 ng/µl as measured with Nanodrop (Thermo Scientific). A crRNA (grl-26_3’_crRNA TTGCGATATAACTTTACGAA) was purchased from IDT, resuspended in IDT duplex buffer to a concentration of 200 µM and annealed to the IDT tracerRNA (200 µM) at 95 °C for 5 minutes. The concentration of the guide RNA was adjusted with water to 62 µM. 0.5 µl of IDT Cas9 protein was mixed with 0.5 µl of guide RNA and incubated for 5 minutes at room temperature. The HRT was heated to 95 °C for 5 minutes and immediately placed on ice. An injection mix was made by adding to 1 µl RNP complex 1 µl 1xTE buffer (pH8), 2 µl HRT and 6 µl water. The mixture was loaded into needles and injected into both gonads of eight young hermaphrodites. No coCRISPR reagents were used. Twenty-four L4 stage progeny of the injected hermaphrodites were picked onto individual plates and kept at 25 °C until they had laid several eggs. Then, each F1 hermaphrodite was picked into 10 µl of proteinase K lysis buffer (Fay & Bender, 2006), frozen in liquid nitrogen, thawed and refrozen 3 times and incubated for 90 minutes at 60 °C and 15 minutes at 95 °C. This worm lysis was used for PCR with primers flanking the insertion site (grl26mKate_chk_F= 5-ACGAACAAGTTGTGGATTCAG-3; grl26mKate_chk_R= 5-GGTTAACTACACCGCTGCC -3).

L4 hermaphrodite progeny of one worm with edits were picked onto individual plates and allowed to lay eggs. Lysis and PCR were performed to identify a homozygous animal. Male progeny of this hermaphrodite showed bright red fluorescence in the tail tip.

### 4.5 Strains used in this study

**Table 1:** Strains used in this study

| designation | genotype | notes |
| --- | --- | --- |
| DF404 | *him-5(e1490) dmd-3(ny21[dmd-3>gfp::3xFLAG::dmd-3]) V; dlg-1(mib23[dlg-1::mCherry-LoxP]) X* | marks DMD-3 and AJs |
| DF405 | *grl-26(ny33[grl-26::mKate]) IV; him-5(e1490) V; qyls10[lam-1p::lam-1::GFP + unc-119(+)]* | marks tai tip cells and BM |
| NF248 | *qyls10[lam-1p::lam-1::GFP + unc-119(+)]* | marks BM |
| DF335 | *nySi1[lin-44>mCherry::PHdomain::lin-44 3'UTR unc-119(+)]II; unc-119III; him-5(e1490)V* | marks tail tip membrane |
| DF288 | *dmd-3 (ny21[GFP::3xFLAG::dmd-3]) him-5(e1490) V* | GFP::DMD-3 |
| DF380 | *grl-26(ny33[grl-26::mKate]) IV; him-5(e1490) V* | GRL-26::mKate |
| DF333 | *unc-119; him-5; cgEx198 [(pJC14) bli-1::GFP + unc-119(+)]* | BLI-1::GFP |
| DF333 | *unc-119; him-5; cgEx198 [(pJC14) bli-1::GFP + unc-119(+)]* | BLI-1::GFP |
| ML2577 | *noah-1(mc68[noah-1::mCh(int)] I; noah-2(mc93[noah-2::flag::gfp(int)]) IV* | NOAH-1::mCherry, NOAH-1::GFP |
| BOX235 | *dlg-1(mib23[dlg-1::mCherry-LoxP]) X* | Marks adherens junctions |

## Author contributions

Karin Kiontke: Conceptualization (equal), investigation (equal), writing - original draft (lead). Porfirio Fernandez: investigation (equal), writing – review and editing (supporting)

Alyssa Woronik: investigation (supporting), writing – review and editing (supporting).

David Fitch: Conceptualization (equal), funding acquisition (lead), writing – review and editing (supporting).

## Acknowledgements

We thank Alison Frand, Michel Labouesse Jeremy Nance and Meera Sundaram for generously sharing fluorescent reporter strains with us. Some strains were provided by the CGC, which is funded by NIH Office of Research Infrastructure Programs (P40 OD010440).

## Funding information

This work was supported by NSF grant 1656736 and NIH grant R01GM141395 to David H. A. Fitch.

## Notes

### Competing Interest Statement

The authors have declared no competing interest.

### Summary of Updates

Author list and references were updated.

## References

1. Buechner, M., Yang, Z., & Al-Hashimi, H. (2020). A Series of tubes: The *C. elegans* excretory canal cell as a model for tubule development. Journal of Developmental Biology, 8(3), 17. 10.3390/jdb8030017

2. Del Rio-Albrechtsen, T., Kiontke, K., Chiou, S.-Y., & Fitch, D. H. A. (2006). Novel gain-of-function alleles demonstrate a role for the heterochronic gene *lin-41* in *C. elegans* male tail tip morphogenesis. Developmental Biology, 297(1), 74–86. 10.1016/j.ydbio.2006.04.472

3. Emmons, S. W. (2005). Male development. WormBook: The Online Review of C. Elegans Biology, 1–22. 10.1895/wormbook.1.33.1

4. Fay, D., & Bender, A. (2006). Genetic mapping and manipulation: Chapter 4--SNPs: introduction and two-point mapping. WormBook: The Online Review of C. Elegans Biology, 1–7. 10.1895/wormbook.1.93.1

5. Geisler, F., Remmelzwaal, S., Jankowski, V., Schmidt, R., Boxem, M., & Leube, R. E. (2023). Intermediate filament network perturbation in the *C. elegans* intestine causes systemic dysfunctions. eLife, 12, e82333. 10.7554/eLife.82333

6. Gill, H. K., Cohen, J. D., Ayala-Figueroa, J., Forman-Rubinsky, R., Poggioli, C., Bickard, K., Parry, J. M., Pu, P., Hall, D. H., & Sundaram, M. V. (2016). Integrity of Narrow Epithelial Tubes in the *C. elegans* Excretory System Requires a Transient Luminal Matrix. PLoS Genetics, 12(8), e1006205. 10.1371/journal.pgen.1006205

7. Herrera, R. A., Kiontke, K., & Fitch, D. H. A. (2016). Makorin ortholog LEP-2 regulates LIN-28 stability to promote the juvenile-to-adult transition in *Caenorhabditis elegans*. *Development (Cambridge*, England), 143(5), 799–809. 10.1242/dev.132738

8. Katz, S. S., Barker, T. J., Maul-Newby, H. M., Sparacio, A. P., Nguyen, K. C. Q., Maybrun, C. L., Belfi, A., Cohen, J. D., Hall, D. H., Sundaram, M. V., & Frand, A. R. (2022). A transient apical extracellular matrix relays cytoskeletal patterns to shape permanent acellular ridges on the surface of adult *C. elegans*. PLoS Genetics, 18(8), e1010348. 10.1371/journal.pgen.1010348

9. Kiontke, K. C., Herrera, R. A., Mason, D. A., Patel, Y., Vernooy, S., Woronik, A., & Fitch, D. H. A. (2024). Tissue-specific RNA-seq defines genes governing male tail tip morphogenesis in C. elegans (p. 2024.01.12.575210). bioRxiv. 10.1101/2024.01.12.575210

10. Kiontke, K. C., Herrera, R. A., Vuong, E., Luo, J., Schwarz, E. M., Fitch, D. H. A., & Portman, D. S. (2019). The long non-coding RNA *lep-5* promotes the juvenile-to-adult transition by destabilizing LIN-28. Developmental Cell, 49(4), 542–555.e9. 10.1016/j.devcel.2019.03.003

11. Lažetić, V., & Fay, D. S. (2017). Molting in *C. elegans*. Worm, 6(1), e1330246. 10.1080/21624054.2017.1330246

12. Lints, R., & Hall, D. H. (2005). WormAtlas Male Handbook—Epithelial System—Hypodermis. WormAtlas. 10.3908/wormatlas.2.7

13. Mason, D. A., Rabinowitz, J. S., & Portman, D. S. (2008). *Dmd-3*, a doublesex-related gene regulated by tra-1, governs sex-specific morphogenesis in *C. elegans*. *Development (Cambridge*, England), 135(14), 2373–2382. 10.1242/dev.017046

14. Mok, D. Z. L., Sternberg, P. W., & Inoue, T. (2015). Morphologically defined sub-stages of *C. elegans* vulval development in the fourth larval stage. BMC Developmental Biology, 15, 26. 10.1186/s12861-015-0076-7

15. Molina-García, L., Lloret-Fernández, C., Cook, S. J., Kim, B., Bonnington, R. C., Sammut, M., O’Shea, J. M., Gilbert, S. P., Elliott, D. J., Hall, D. H., Emmons, S. W., Barrios, A., & Poole, R. J. (2020). Direct glia-to-neuron transdifferentiation gives rise to a pair of male-specific neurons that ensure nimble male mating. eLife, 9, e48361. 10.7554/eLife.48361

16. Nelson, M. D., Zhou, E., Kiontke, K., Fradin, H., Maldonado, G., Martin, D., Shah, K., & Fitch, D. H. A. (2011). A Bow-Tie Genetic Architecture for Morphogenesis Suggested by a Genome-Wide RNAi Screen in *Caenorhabditis elegans*. PLOS Genetics, 7(3), e1002010. 10.1371/journal.pgen.1002010

17. Nguyen, C. Q., Hall, D. H., Yang, Y., & Fitch, D. H. (1999). Morphogenesis of the *Caenorhabditis elegans* male tail tip. Developmental Biology, 207(1), 86–106. 10.1006/dbio.1998.9173

18. Remmelzwaal, S., Geisler, F., Stucchi, R., van der Horst, S., Pasolli, M., Kroll, J. R., Jarosinska, O. D., Akhmanova, A., Richardson, C. A., Altelaar, M., Leube, R. E., Ramalho, J. J., & Boxem, M. (2021). BBLN-1 is essential for intermediate filament organization and apical membrane morphology. Current Biology: CB, 31(11), 2334–2346.e9. 10.1016/j.cub.2021.03.069

19. Schindler, A. J., & Sherwood, D. R. (2013). Morphogenesis of the Caenorhabditis elegans vulva. Wiley Interdisciplinary Reviews. Developmental Biology, 2(1), 75–95. 10.1002/wdev.87

20. Serrano-Saiz, E., Oren-Suissa, M., Bayer, E. A., & Hobert, O. (2017). Sexually Dimorphic Differentiation of a C. elegans Hub Neuron Is Cell Autonomously Controlled by a Conserved Transcription Factor. Current Biology: CB, 27(2), 199–209. 10.1016/j.cub.2016.11.045

21. Starr, D. A. (2019). A network of nuclear envelope proteins and cytoskeletal force generators mediates movements of and within nuclei throughout *Caenorhabditis elegans* development. *Experimental Biology and Medicine (Maywood*, N.J.), 244(15), 1323–1332. 10.1177/1535370219871965

22. Stiernagle, T. (2006). Maintenance of C. elegans. WormBook: The Online Review of C. Elegans Biology, 1–11. 10.1895/wormbook.1.101.1

23. Sulston, J. E., Albertson, D. G., & Thomson, J. N. (1980). The *Caenorhabditis elegans* male: Postembryonic development of nongonadal structures. Developmental Biology, 78(2), 542–576. 10.1016/0012-1606(80)90352-8

24. White, J. G., Southgate, E., Thomson, J. N., & Brenner, S. (1986). The structure of the nervous system of the nematode Caenorhabditis elegans. *Philosophical Transactions of the Royal Society of London. Series B*, Biological Sciences, 314(1165), 1–340. 10.1098/rstb.1986.0056

25. Woronik, A., Kiontke, K., Jallad, R. S., Herrera, R. A., & Fitch, D. H. A. (2022). Laser Microdissection for Species-Agnostic Single-Tissue Applications. Journal of Visualized Experiments: JoVE, 181. 10.3791/63666

26. Xiao, Y., Yee, C., Zhao, C. Z., Martinez, M. A. Q., Zhang, W., Shen, K., Matus, D. Q., & Hammell, C. (2023). An expandable FLP-ON::TIR1 system for precise spatiotemporal protein degradation in *Caenorhabditis elegans*. Genetics, 223(4), iyad013. 10.1093/genetics/iyad013

27. Zahreddine, H., Zhang, H., Diogon, M., Nagamatsu, Y., & Labouesse, M. (2010). CRT-1/calreticulin and the E3 ligase EEL-1/HUWE1 control hemidesmosome maturation in *C. elegans* development. Current Biology: CB, 20(4), 322–327. 10.1016/j.cub.2009.12.061

28. Zhao, X., Yang, Y., Fitch, D. H. A., & Herman, M. A. (2002). TLP-1 is an asymmetric cell fate determinant that responds to Wnt signals and controls male tail tip morphogenesis in *C. elegans*. *Development (Cambridge*, England), 129(6), 1497–1508. 10.1242/dev.129.6.1497

